# PhenoMe: An interactive AI framework for interpretable, microscopy-driven phenotyping

**DOI:** 10.64898/2026.10.29.762637

**Authors:** Aitor González-Marfil, Lenka Backová, Renaud Bernatchez, Iván Hidalgo-Cenalmor, Antoine Dufour, Guillaume Jacquemet, Flavie Lavoie-Cardinal, Ignacio Arganda-Carreras, Estibaliz Gómez-de-Mariscal

## Abstract

Extracting meaningful biological patterns from microscopy image datasets requires tools that quantitatively connect visual similarities and experimental context. Deep learning models, and in particular novel foundation models, have the capacity to translate images into high-dimensional embeddings that can better represent complex visual patterns. Using these complex representations analytically and linking them to experimental information, it is possible to build advanced exploratory and phenotyping capabilities. Here, we present PhenoMe, an open-source, model-agnostic framework to handle pretrained foundation model-derived image representations alongside quantitative image properties and experimental metadata. PhenoMe integrates multiple scriptable and interactive workflows for embedding space visualisation, dimensionality reduction and clustering, similarity-based comparison and retrieval of representative images for each phenotype, among others. It also supports correlation analyses that link the embedding structure to interpretable image properties. We demonstrate PhenoMe’s capabilities on seven public microscopy datasets, ranging from cell painting assays, nanoscale synaptic images, drug-response analyses and dynamic cell-phenotyping. These case studies show how readily available image representations can support data contextualisation, curation and analysis. Through guided notebooks, reproducible configurations, extensible model wrappers and a napari plugin, PhenoMe makes representation-based microscopy exploration accessible to researchers without dedicated technical expertise. PhenoMe lays the groundwork for applying foundation models and representation learning to microscopy data analysis.

## Introduction

Microscopy images contain a wealth of biological information, but extracting biological insight from increasingly large and complex image collections remains challenging. Manual inspection is time-consuming and difficult to scale, whereas conventional quantitative analyses rely on predefined measurements that may capture only part of the visual variation in the data. Representation-learning methods offer a complementary approach within AI-driven image analysis by mapping each image or image object into a high-dimensional vector of extracted visual features, known as an embedding. A key benefit is that across a dataset, the computed image embeddings define a dense latent feature space in which images can be organised, compared and related to experimental conditions. Such embeddings can be obtained from vision models trained for different objectives, including classification, object detection, segmentation and prediction of biological responses (1–6). Accordingly, AI-derived embeddings are not predefined image features but learnt representations shaped by the vision model’s architecture, training data, learning objective, and training approach.

Self-supervised learning (SSL) approaches are particularly interesting because the model’s learnt visual features are derived from the data rather than from human-assigned image labels. That is, the resulting embeddings may capture variations that are not constrained by a predefined supervised task. Although such representations are not inherently interpretable, their latent structure can reveal patterns that analyses based solely on predefined features may overlook. Previous works based on SSL approaches have indeed captured single-cell phenotypic variation and mechanisms of action (7, 8), identified extracellular-matrix-dependent morphology (9), and recovered morphodynamic states from live-cell imaging (10). More recent studies have extended these approaches to large morphological-profiling datasets (11, 12), developed interpretable representations of cellular morphology (13), and related image-derived phenotypes to genetic fingerprints, perturbations, neuronal activity and spatial transcriptomics (14–18). Collectively, these studies demonstrate that learnt feature spaces can support the organisation of cellular images, the characterisation of phenotypic variation, the association of phenotypes with molecular and experimental variables, and the discovery of predictive patterns (19).

Vision foundation models extend this strategy by applying SSL at scale to learn representations from large, diverse datasets that can be reused across downstream applications (20). Consequently, they can serve as feature extractors without task-specific retraining, although their transferability depends on the alignment between the source data and target domain. General-purpose models such as DINOv2 (21) have been applied across microscopy modalities (22), while microscopy-oriented foundation models increasingly target cellular morphology, protein localisation and perturbation responses. Examples include Cell-DINO (1), the proteome-aware SubCell models (23), and a foundation model for stimulated-emission-depletion microscopy (STED-FM) (24), whose embeddings can be used for various downstream tasks, including clustering, image retrieval, segmentation and generation of STED images of various neuronal structures such as the F-Actin cytoskeleton and synaptic protein clusters. These developments expand the availability of reusable image representations, but they do not by themselves provide the analytical workflows required to explore and interpret those representations in a biological context.

A practical gap therefore remains between generating image embeddings and using them for biological analysis. Researchers must extract and store embeddings, link them to experimental metadata and source images, examine the structure of the resulting latent space, and determine which measurable image properties or experimental variables explain that structure. Existing software addresses complementary parts of this workflow. CellProfiler provides established pipelines for image processing, object segmentation, and the quantification of predefined features (25), whereas FiftyOne supports the interactive exploration of images, labels, and embeddings in general computer-vision applications (26). An accessible, model-agnostic system that connects learnt representations with microscopy-specific measurements, source images and experimental variables within the Python data-analysis ecosystem would make this workflow more reproducible, interpretable and extensible.

Here, we present **PhenoMe**, an open-source framework for the interactive exploration, contextualisation and analysis of microscopy images using learnt representations (**Fig. 2**). PhenoMe provides a bridging software layer between pretrained vision models and microscopy image analysis. It connects embeddings with source images, experimental metadata and classical image measurements, enabling users to inspect latent-space structure, compare experimental conditions and investigate the measurable properties associated with observed patterns. We demonstrate the workflow following a common interpretive sequence: we first identified structure in the learnt representation, related that structure to experimental metadata, retrieved the corresponding images for visual assessment and finally tested its association with measurable image properties. This sequence is applied to seven publicly available datasets that include human and bacterial cells, nanoscale synaptic structures, compound profiling, dose-response experiments and cell dynamics.

## Results

### PhenoMe framework – integrated learnt representations, image properties and metadata

PhenoMe is a modular, model-agnostic Python framework that combines deep-learning embeddings, quantitative image properties and experimental metadata for exploratory microscopy analysis (**Fig. 1**). It provides a scriptable Python API, guided Jupyter notebooks that can run locally or through Google Colab and a user-friendly interface within napari (**Supplementary Fig. 6**). The input pipeline supports 2D microscopy images, including multichannel, paired images with segmentation masks or time-lapse sequences, spanning a wide range of microscopy modalities. Multispectral image embeddings are computed by converting the images to RGB before encoding or by encoding each channel independently and concatenating the resulting embeddings. When segmentation masks are provided, images are cropped using the masks, creating one image per object. Time-lapse image data can be integrated by splitting each video into 2D images and including the corresponding time point reference within the metadata file: a CSV text file that maps each image to additional features as for example, a replicate, treatment, cell type, or acquisition time point. This structure provides a standardised mechanism for associating experimental metadata with image collections and integrating additional features from complementary analytical software. This flexibility also enables PhenoMe to accommodate large image collections with heterogeneous organisational structures. Further technical information is available in the Supplementary Material section C. PhenoMe integrates several ready-to-use pretrained models, specifically the small, base, large, and giant variants of DINOv2, with a model wrapper that supports the addition of custom models. Alongside image embeddings, PhenoMe extracts intensity measurements, grey-level cooccurrence matrix (GLCM) texture features, and imagequality metrics. When masks are available, it also calculates foreground and background properties, concentric-ring features and object-shape descriptors. Embeddings, properties and metadata are stored internally, supporting joint dimensionality-reduction (e.g., t-SNE, UMAP, and principal component analysis (PCA)), clustering, correlation analysis and further statistical analysis. The interactive visualiser links the AI-derived feature space to the source images and lets users filter, select, and assign provisional labels (**Fig. 2**). Although PhenoMe can run completely on a CPU-based system, GPU acceleration is recommended for AI-derived embedding extraction. The framework is released as an MIT-licensed package with documentation, tutorials and reproducible examples for its broad adoption and integration within the community.

**Fig. 1.**
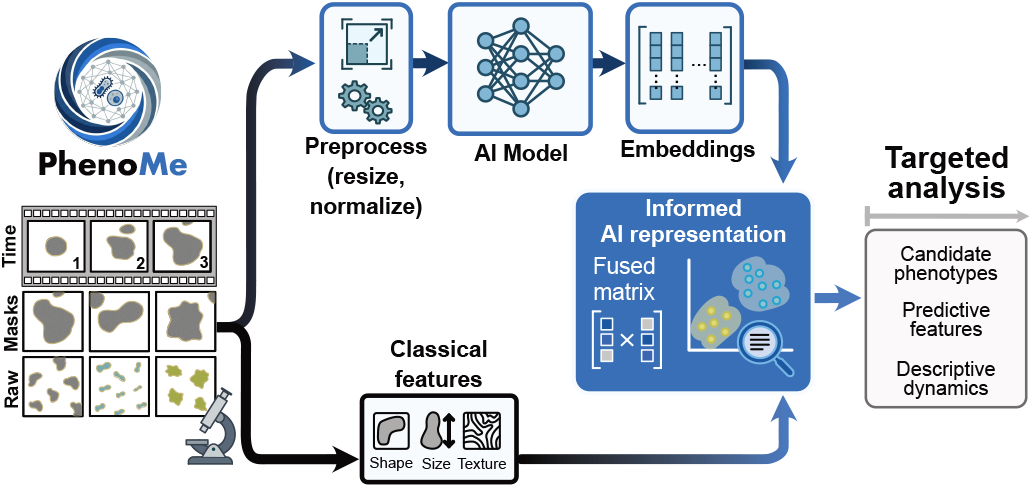
PhenoMe framework for interactive microscopy data exploration. PhenoMe processes informed 2D image collections (e.g., multichannel, paired raw and segmentation images, and time-lapse acquisitions) and combines embeddings from pretrained models with classical morphological, intensity, texture and quality measurements, as well as user-provided experimental metadata. The integrated representation supports dimensionality reduction, clustering, correlation analysis and interactive inspection. The resulting embeddings, selected observations, quality flags and clusters or provisional labels can be exported for subsequent dedicated quantitative analyses.

**Fig. 2.**
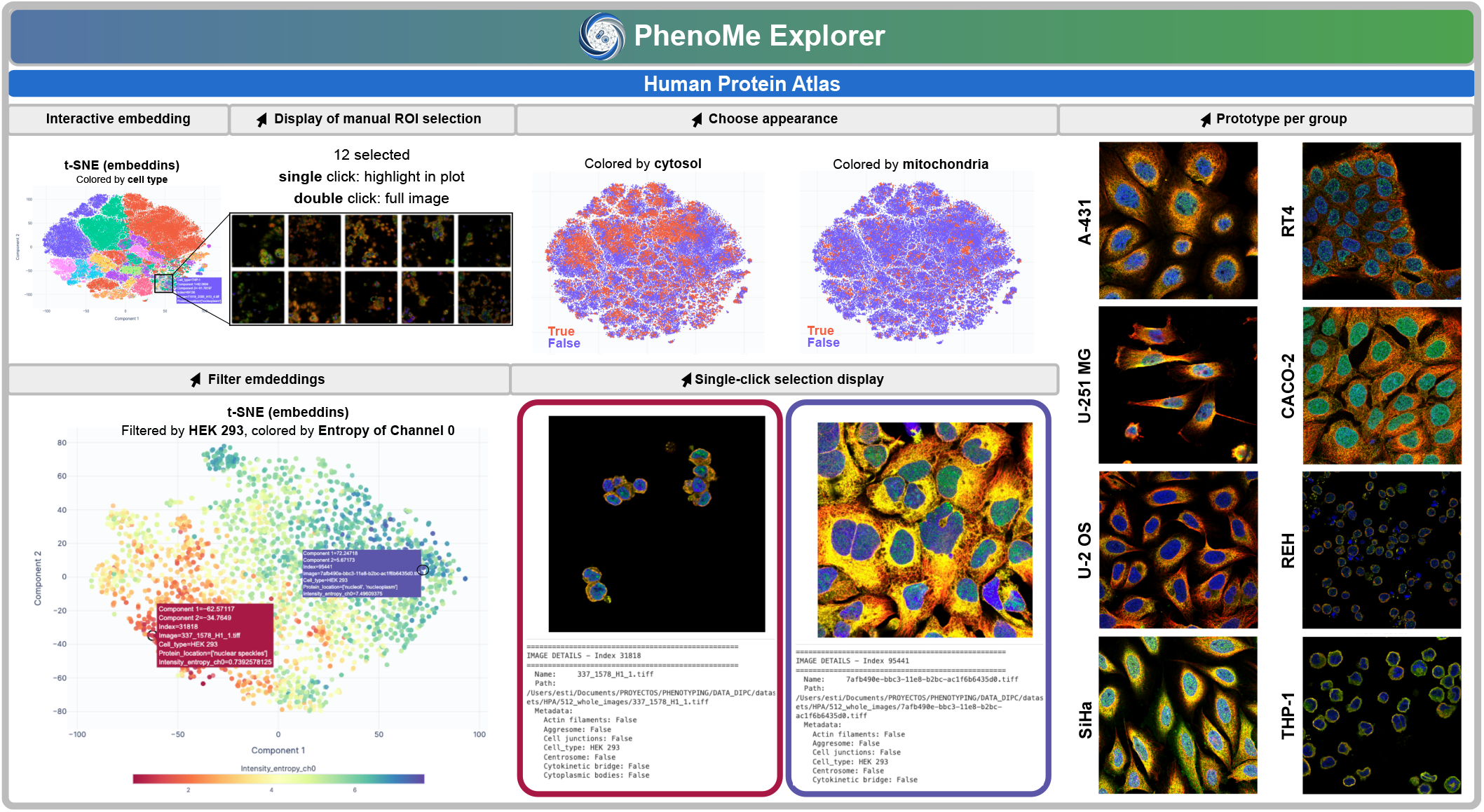
PhenoMe’s graphical interface. (Interactive embedding) PhenoMe supports interaction with the image representations derived from pre-trained models: it allows inspecting the data distribution, colouring and filtering the representations using different variables (choose appearance and filter embeddings), as well as retrieving images from data points (display of manual ROI selection and single-click selection display) and representatives of each cluster (prototype per group), along with their metadata. This allows linking locations in the representation space to images, measurements and metadata. Image data clusters can be obtained from the embeddings using cluster analysis or can be pre-established by a given categorical variable, as for example the cell type in this case. Supplementary Fig. 7 shows representative candidates for all cell types. The example here displays the t-SNE projections of immunofluorescence images from the Human Protein Atlas embedded with Cell-DINO.

### Comparison of embedding spaces

PhenoMe provides a model-agnostic wrapper that enables the use of different state-of-the-art models, including domain-specific foundation models. This flexibility helps users identify model representations that are suited to a particular biological question and to examine how different models encode biological variation. It also facilitates the integration and benchmarking of newly developed architectures, ensuring that PhenoMe remains adaptable to new network architectures and domainspecific models.

We embedded immunofluorescence images from the Human Protein Atlas (HPA) (28) using Cell-DINO (1), DINOv2 ViTS/14 (21) and ConvNeXt Large (29) (**Fig. 3a**). Interestingly, in all three embedding spaces, the resulting t-SNE projections revealed a cell-type-dependent organisation, with morphologically similar cell types located close to one another or partially overlapping (**Fig. 3a**). Such visualisations facilitate the rapid identification of phenotypes from different categories that occupy neighbouring regions of the latent space, thereby supporting the interpretation of shared image patterns and morphological characteristics. At the same time, cell type groups’ topology and degree of separation varied substantially among models. DINOv2 produced compact, well-defined clusters for several abundant cell types, whereas Cell-DINO retained a distinct structure for some smaller cell type groups. By comparison, the ConvNeXt embedding exhibited greater overlap among multiple cell types. These differences likely reflect variation in model architecture, training data, and training strategies. Thus, the capacity to easily load, explore and compare model embeddings helps in choosing an adequate representation of the data, but also in studying deeper AI learning capabilities.

**Fig. 3.**
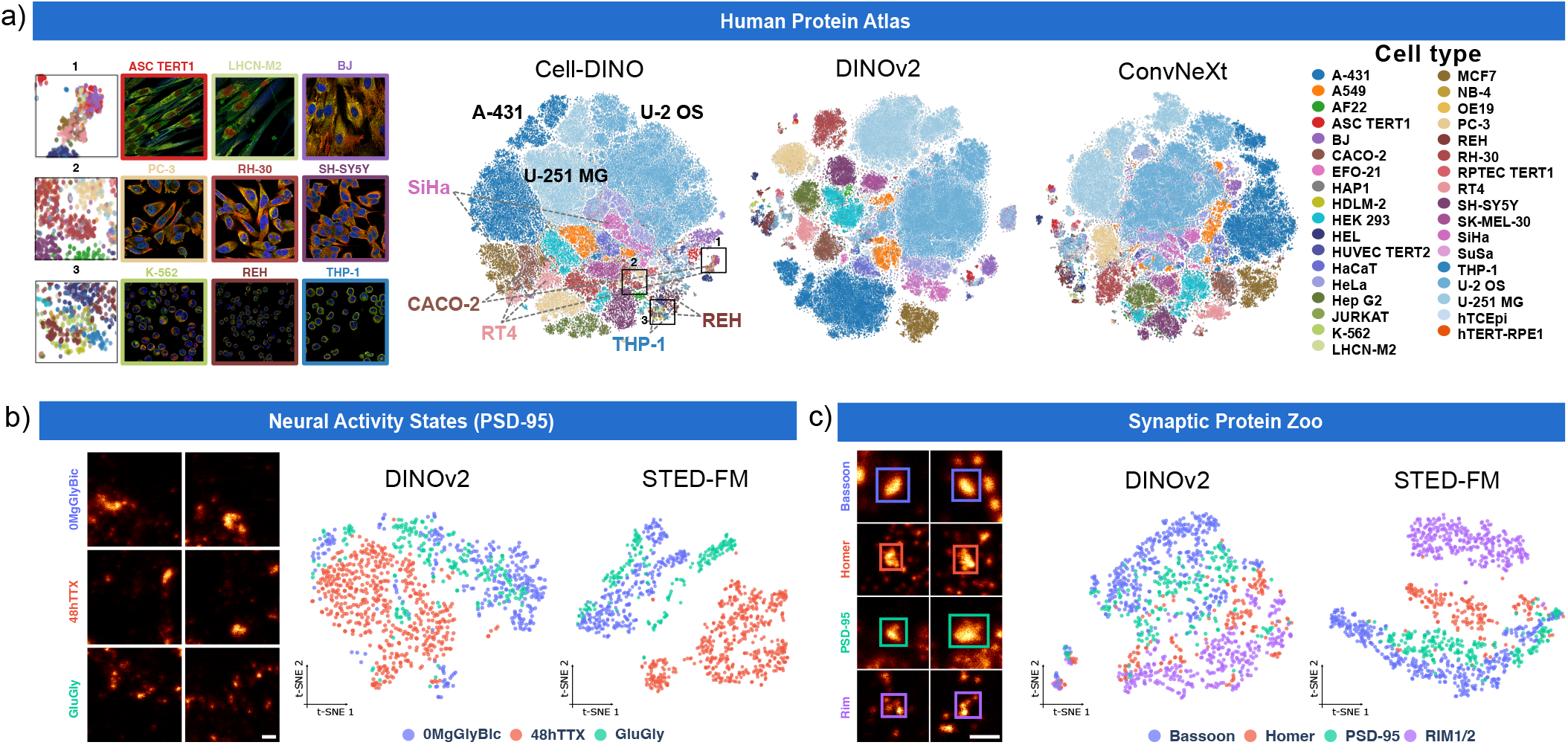
Universal framework for foundation models. a) Example images of the Human Protein Atlas immunofluorescence dataset together with t-SNE projections of their embeddings using Cell-DINO, DINOv2 ViT-S/14, or ConvNeXt Large. Points are coloured by cell type; highlighted cell types in Cell-DINO’s representation correspond to the visual examples in Fig. 2. Supplementary Fig. 7 shows representative candidates for all cell types. b) Representative STED images of PSD-95 from neurons subjected to 0 Mg^2+^/Gly/Bic, 48 h TTX, or Glu/Gly treatment, together with t-SNE projections of their DINOv2 and STED-FM embeddings. Points are coloured by experimental condition. c) Representative clusters of Bassoon, Homer1c, PSD-95, and RIM1/2 from the Synaptic Protein Zoo dataset, together with t-SNE projections of their DINOv2 and STED-FM embeddings. b-c) Scale bars = 500 nm.

We also compared the representations obtained by domain-specific models against DINOv2. For this, we used two different image datasets of neural synaptic protein clusters acquired with STED microscopy and STED-FM, a foundation model trained specifically on STED microscopy images (24) (**Fig. 3b–c**). The Neural Activity States dataset contains STED images of the postsynaptic scaffold protein PSD-95 in fixed hippocampal neurons subjected to three activity-modulating treatments (24, 30): 48*h* chronic treatment with the sodium-channel blocker tetrodotoxin (TTX), which induces homeostatic synaptic upscaling; 0 Mg^2+^/Gly/Bic stimulation, which induces chemical long-term potentiation (cLTP); and Glu/Gly stimulation, which induces chemical long-term depression (cLTD). Although the DINOv2 embedding captured some treatment-associated structure, the three conditions remained substantially intermixed. In the STED-FM embedding, the 48*h* TTX samples occupied more distinct regions, while the two acute stimulation conditions formed partially separated clusters (**Fig. 3b**). Thus, the representation learnt through domain-specific pretraining appeared to be more closely aligned with the experimentally induced activity states.

In addition, we analysed images from the Synaptic Protein Zoo dataset (24, 31)^1^. Individual synaptic protein clusters were extracted using their corresponding segmentation masks and labelled according to protein identity:

Bassoon, Homer1c, PSD-95, or RIM1/2. Protein classes were broadly distributed and partially intermixed in the DI-NOv2 embedding. In contrast, the STED-FM representation showed stronger organisation by protein identity, most notably through the formation of a distinct RIM1/2 region and more coherent groups for the remaining proteins (**Fig. 3c**). Together, these examples show that the biological structure captured by an embedding depends strongly on the selected backbone and its pretraining domain. By enabling different representations to be loaded and explored through a common interface, PhenoMe supports both the selection of models for downstream phenotyping and the investigation of how general-purpose and domain-specific models encode biological information.

### Interactive contextualisation reveals phenotypes and acquisition artefacts

To showcase PhenoMe’s capabilities in microscopy-driven research, we next analysed two datasets from the Broad Bioimage Benchmark Collection (BBBC) (32). BBBC021v1 contains multiplexed fluorescence images of Michigan Cancer Foundation-7 (MCF7) cells (human breast adenocarcinoma cell line) exposed to targeted and cancer-relevant cytotoxic compounds with annotated mechanisms of action (MoAs) (33). A t-SNE projection of DINOv2 ViT-g/14 embeddings was substantially organised by MoA (**Fig. 4a**). The interactive selection of microtubule-destabilisers revealed a compound-level phenotypic heterogeneity within this MoA (**Fig. 4b**). Colchicine formed a separate group, nocodazole occupied a partially distinct region, and vincristine and demecolcine overlapped extensively. To relate the projection to measurable properties and identify predictive features for these phenotypes, we used PhenoMe’s integrated Pearson correlation analysis (34) between the first two t-SNE coordinates (35) and classical features (**Fig. 4c**). The first coordinate is strongly correlated with the entropy, mean and minimum intensity, standard deviation and sharpness of the *β*-tubulin channel. This aligns with the visual differences between colchicine and the remaining conditions, with microtubules displaying a fibrillary organisation when cells are treated with colchicine (**Fig. 4d**). In contrast, the second coordinate is strongly correlated with DNA sharpness, aligning with the observed decomposed nuclei when cells are treated with demecolcine and vincristine.

**Fig. 4.**
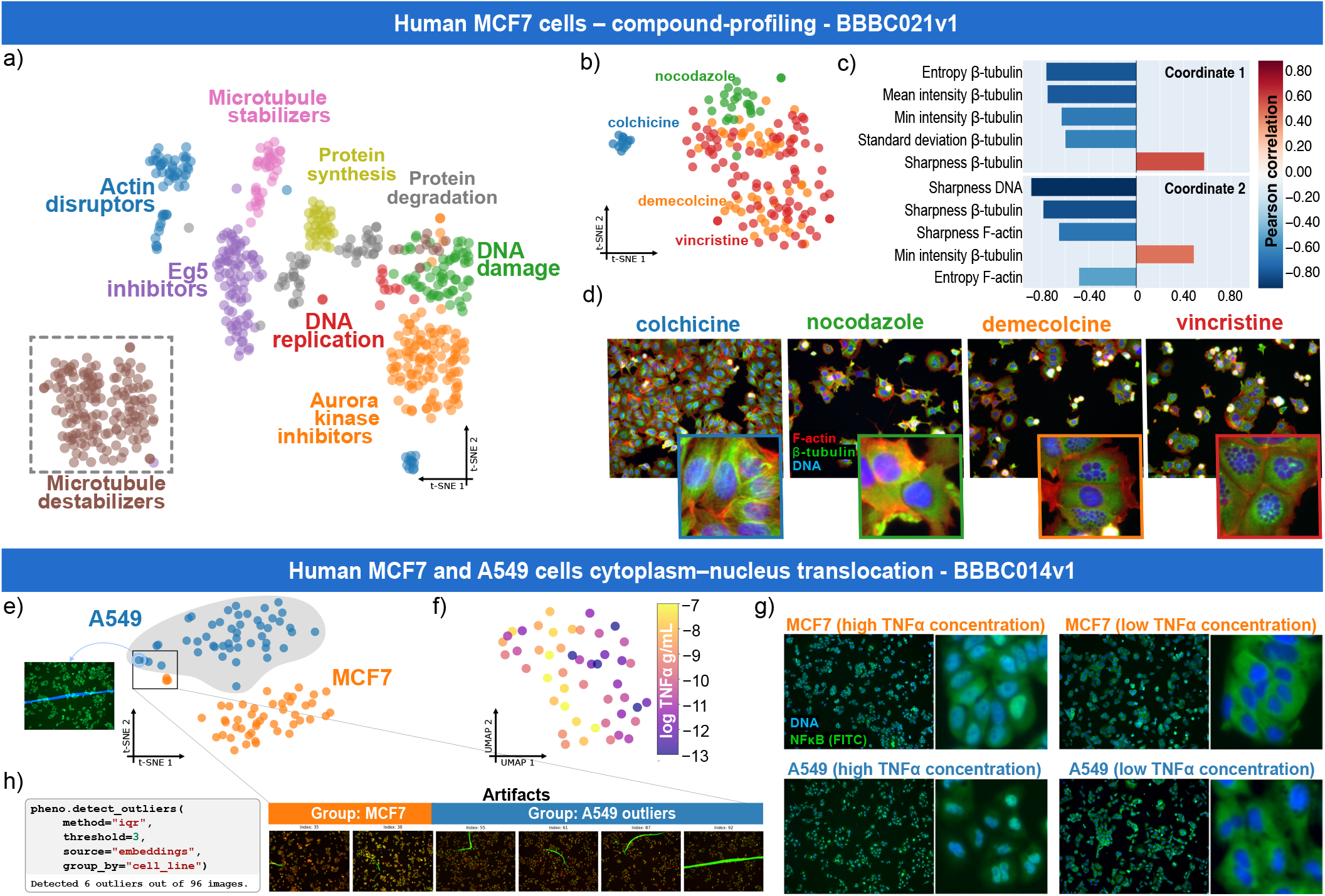
Representation-based data exploration and contextualisation. a) t-SNE projection of DINOv2 ViT-g/14 embeddings from a subset of the BBBC021v1 compound-profiling dataset comprising MCF7 cells treated with 19 compounds (27). Points are coloured by annotated mechanism of action (MoA); the dashed box identifies microtubule-destabilising compounds. b) t-SNE projection of microtubule-destabiliser embeddings, coloured by compound. Colchicine and nocodazole have relatively distinct distributions, whereas vincristine and demecolcine overlap substantially. c) Pearson correlation coefficients between each t-SNE coordinate and classical image properties for the filtered dataset in b). d) Representative multichannel fluorescence images for each microtubule-destabilising compound. Channels correspond to F-actin (red), *β*-tubulin (green) and DNA (blue). e) t-SNE projection of DINOv2 ViT-g/14 embeddings from BBBC014v1, coloured by cell type. f) UMAP projection of the A549 subset, coloured by log_10_ TNF-*α* concentration in g mL^*−*1^. The first coordinate is associated with concentration (Spearman *ρ* = *−*0.53). g) Representative MCF7 and A549 images under high and low TNF-*α* conditions (log_10_ concentration values of *−*7 and *−*11 in g mL^*−*1^, respectively), illustrating positive and negative NF-*κ*B cytoplasm-to-nucleus translocation. Nuclei are labelled with DAPI (blue), and NF-*κ*B is detected in the FITC channel (green). h) Boxed outliers in e) contain acquisition artefacts that can be manually inspected individually or identified by running the outlier detector of PhenoMe.

Thus, with PhenoMe’s functionalities and capacity to analytically handle embeddings, one can easily identify and link the phenotypic heterogeneity among compounds to separable quantitative image properties, such as microtubule organisation and nuclear integrity in this case.

BBBC014v1 contains dual-channel images–DAPI and FITC for the nucleus and cytoplasm, respectively–of MCF7 and A549 alveolar epithelial cells exposed to 12 concentrations of tumour necrosis factor alpha (TNF-*α*). TNF-*α* induces NF-*κ*B translocation from the cytoplasm to the nucleus, changing the spatial relationship between FITC and DAPI signals. A t-SNE projection of DINOv2 ViT-g/14 embeddings separated most observations into two groups, matching with the two cell lines (**Fig. 4e)**. When inspecting each cell type individually, within the A549 subset, the first coordinate of a UMAP projection was moderately associated with TNF-*α* concentration (Spearman *ρ* = *−*0.53; **Fig. 4f**) and with the image-level standard deviation of the FITC signal (Pearson *r* = *−*0.62). Representative images showed the corresponding concentration-dependent redistribution of NF-*κ*B (**Fig. 4g)**. In BBBC014v1, several images affected by acquisition artefacts can be quickly identified either individually through PhenoMe’s interactive image retrieval or automatically using its outlier-detection functionality (**Fig. 4h**). This outlier identification and inspection supports faster quality control before downstream analysis. Similarly, linked metadata, images, and quantitative measurements help disentangle variation in the embedding space by revealing whether it reflects biological factors, technical factors, or both.

### Trajectories in the embedding space capture continuous phenotypic variation

Next, we showcase how PhenoMe could support the exploration of time-resolved phenotypic responses and dynamic biological processes. Namely, labelling and tracking changes in the representation space allows pattern-based delineation of dynamic phenotypic changes, such as the onset and progression of treatment-associated phenotypes. We first used a confocal microscopy dataset of fixed *Escherichia coli* (E. coli) cells stained for DNA and membrane and expressing MreB-sfGFP^sw^ (36).

Cultures were exposed to antibiotic perturbations and fixed after 0, 2, 5, 10, 20, 30, and 60 min. Instance-segmentation masks supplied with the dataset enabled PhenoMe to integrate conventional single-cell morphometric features. Seeing the projection of DINOv2 ViT-g / 14 t-SNE at different time points revealed a treatment-associated dynamic redistribution of the data (**Fig. 5a** and Supplementary **Fig. 8a**). At 0 min, most conditions overlap, and the features showing the strongest Pearson correlations describe bacterial morphology and correlate positively with the first coordinate. Visual embedding inspection suggests some fine patterns related to bacterial division and elongation (Supplementary **Fig. 8c**). Early cell responses to treatment can be distinguished after 20 mins of treatment exposure and explained by changes in the spatial distribution of the three channels, particularly DNA channel (**Fig. 5b** and Supplementary **Fig. 8d**). At 60 min, a treatment-associated organisation, with related cellular morphologies occupying neighbouring or overlapping regions, is revealed (**Fig. 5a**). Namely, feature correlation analysis results in high correlations with both t-SNE coordinates and identified minor-axis length as the property most strongly associated with the first t-SNE coordinate (Pearson *ρ ≈−*0.81; **Fig. 5b**). Recolouring the projection by this property revealed a gradient in bacterial width and rounding (**Fig. 5c** and Supplementary **Fig. 8e**). Using PhenoMe, we calculated the distance between the mean embedding for each condition and time point and that of the control at 0 min. The resulting treatment-response trajectories revealed early responses to rifampicin and chloramphenicol, pronounced late responses to mecillinam and MP265, and only a small shift following sodium azide treatment (**Fig. 5d**), consistent with the treatment-dependent dynamics reported in the source study (36).

**Fig. 5.**
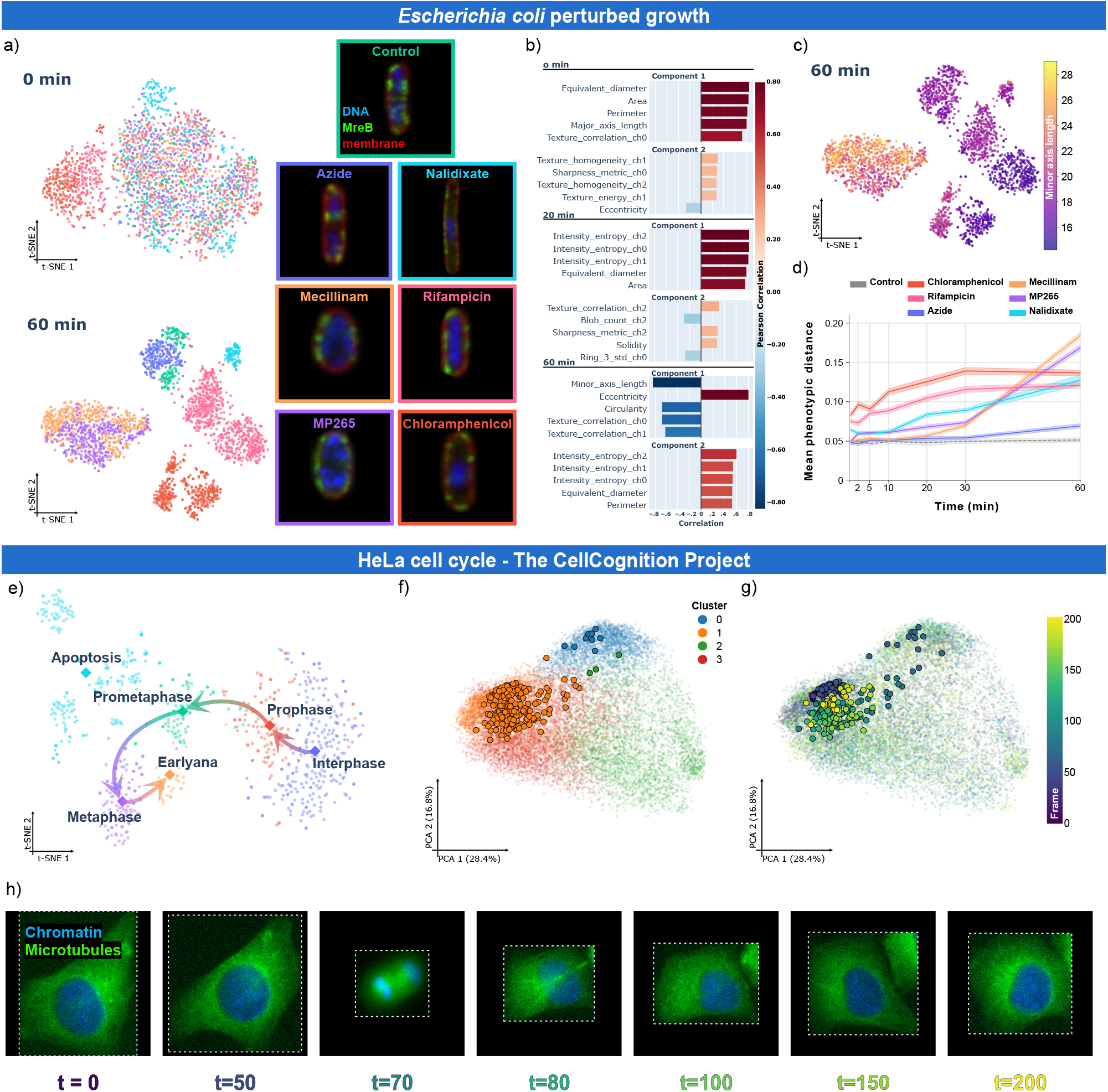
Dynamic phenotyping and predictive continuous biological variation. a) t-SNE projection of DINOv2 ViT-g/14 embeddings from *E. coli* cells at 0 and after 60 min of treatment, coloured by perturbation. Images retrieved with PhenoMe illustrate treatment-associated morphologies after 60 min of treatment. b) Pearson correlation analyses relating the embedding projection at 0, 20 and 60 min time points to classical properties. c) The same projection coloured by minor-axis length (in pixels), an indicator of bacterial width and rounding. Minor-axis length was strongly associated with the first t-SNE coordinate (Pearson *r ≈−*0.81). d) Mean representation space distance of each treatment group from the centroid of the 0-min control group over time. Shading denotes 95% confidence intervals. e) t-SNE projection of DINOv2 ViT-g/14 embeddings from the CellCognition HeLa dataset, coloured by annotated cell-cycle phase (interphase, prophase, prometaphase, metaphase, early anaphase, and apoptosis). The embeddings form a continuous cell-cycle-associated trajectory, whereas apoptosis is comparatively separated. f-g) PCA projection of embedded single-cell crops from a time-lapse acquisition of dividing HeLa cells, coloured by f) predicted clusters and g) frame number. Highlighted points form the single-cell track shown in h).

**Fig. 6.**
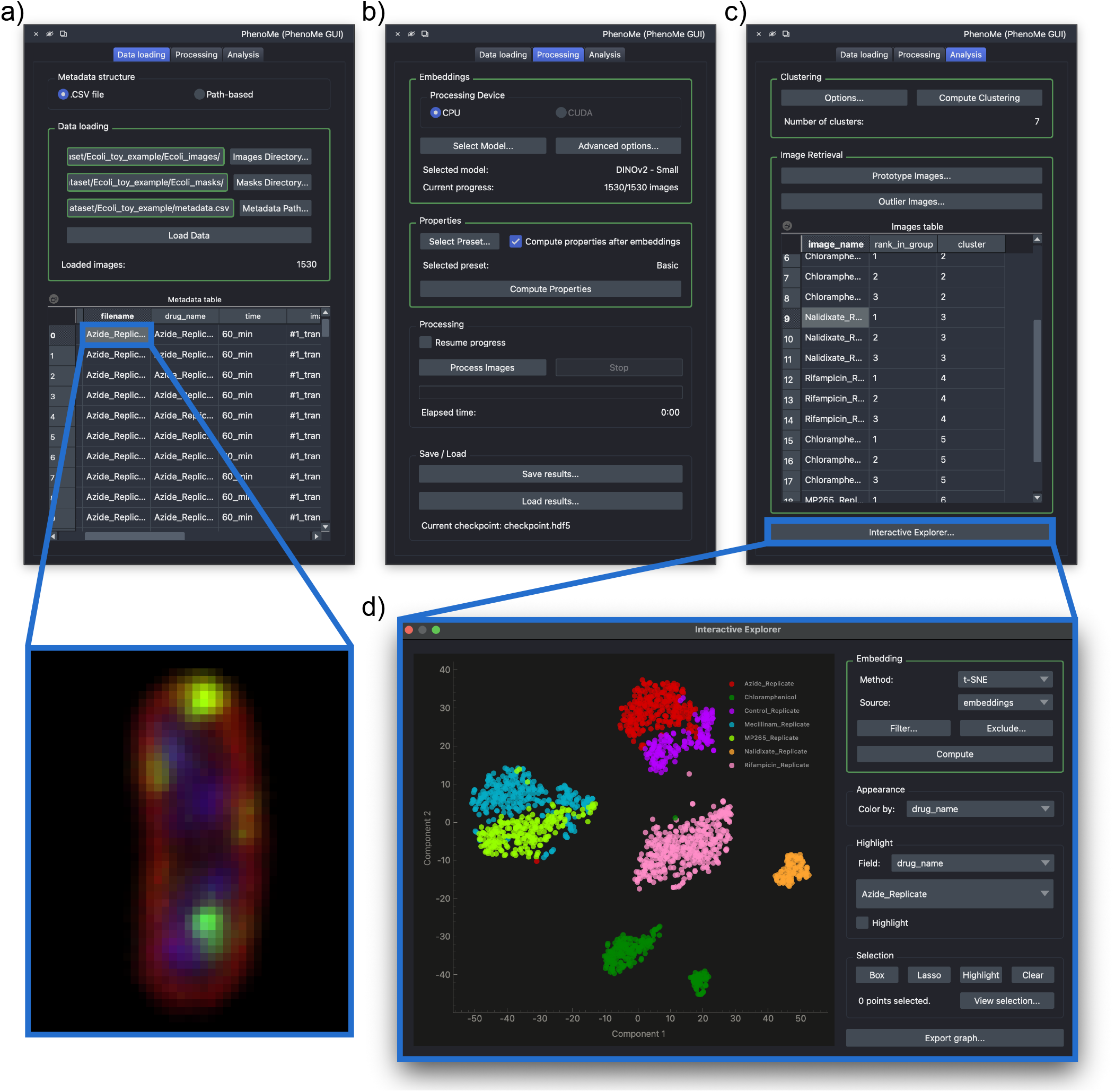
PhenoMe plugin for the napari image viewer. a) Data loading panel, allowing users to import images, masks, and metadata from specified paths. Selecting an image from the data table displays it in the napari viewer. b) Embedding extraction and property calculation panel, allowing users to select from the available foundation models and quantitative properties. c) Results analysis panel, providing options for clustering, retrieving prototype and outlier images based on identified clusters or predefined groups, and visualizing embeddings. d) Interactive explorer for visualizing embeddings using a selected dimensionality reduction algorithm. Filters can be applied to explore data subsets based on metadata or quantitative properties.

**Fig. 7.**
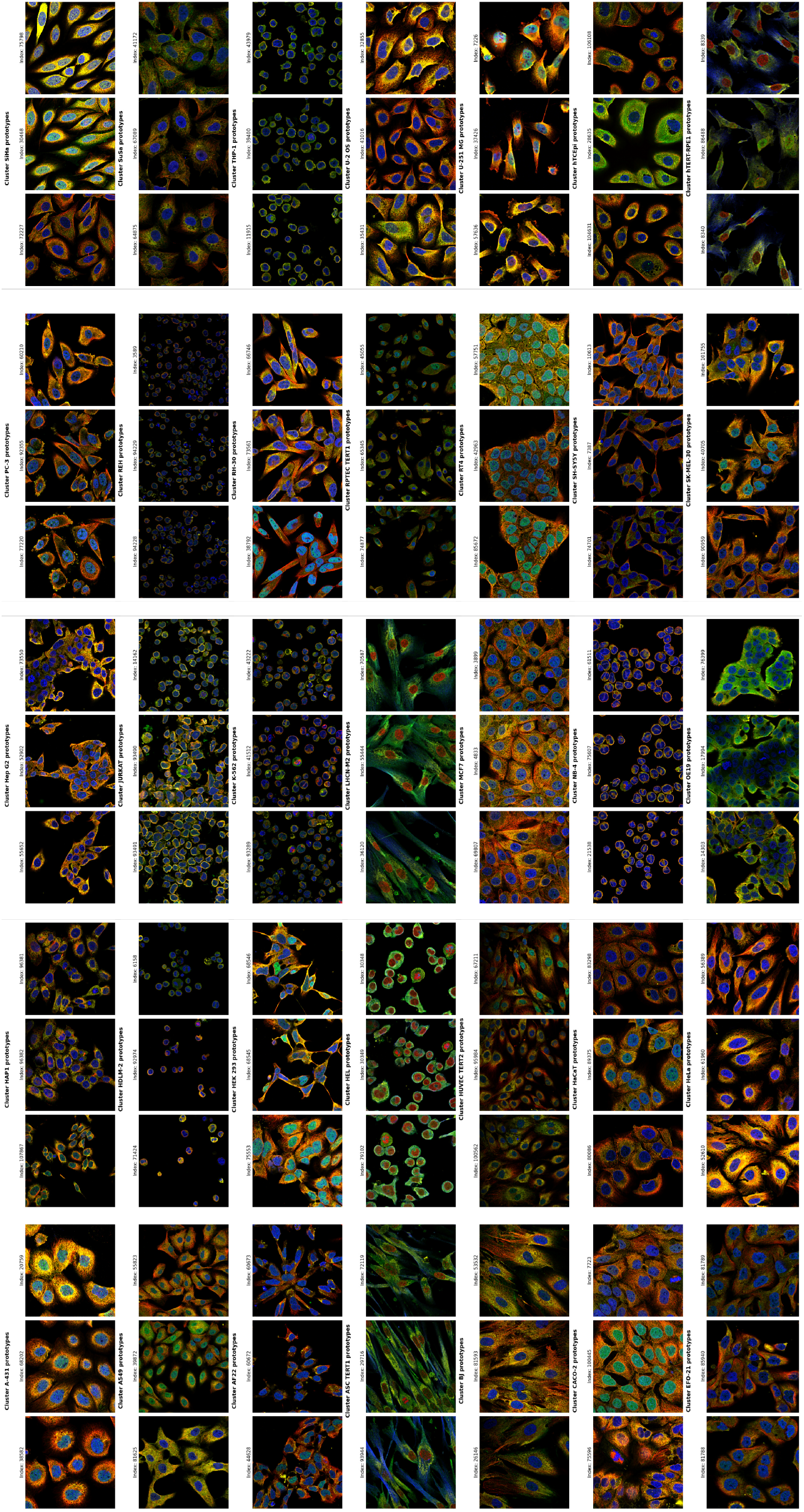
Prototype images of the HPA dataset per cell type cluster.

**Fig. 8.**
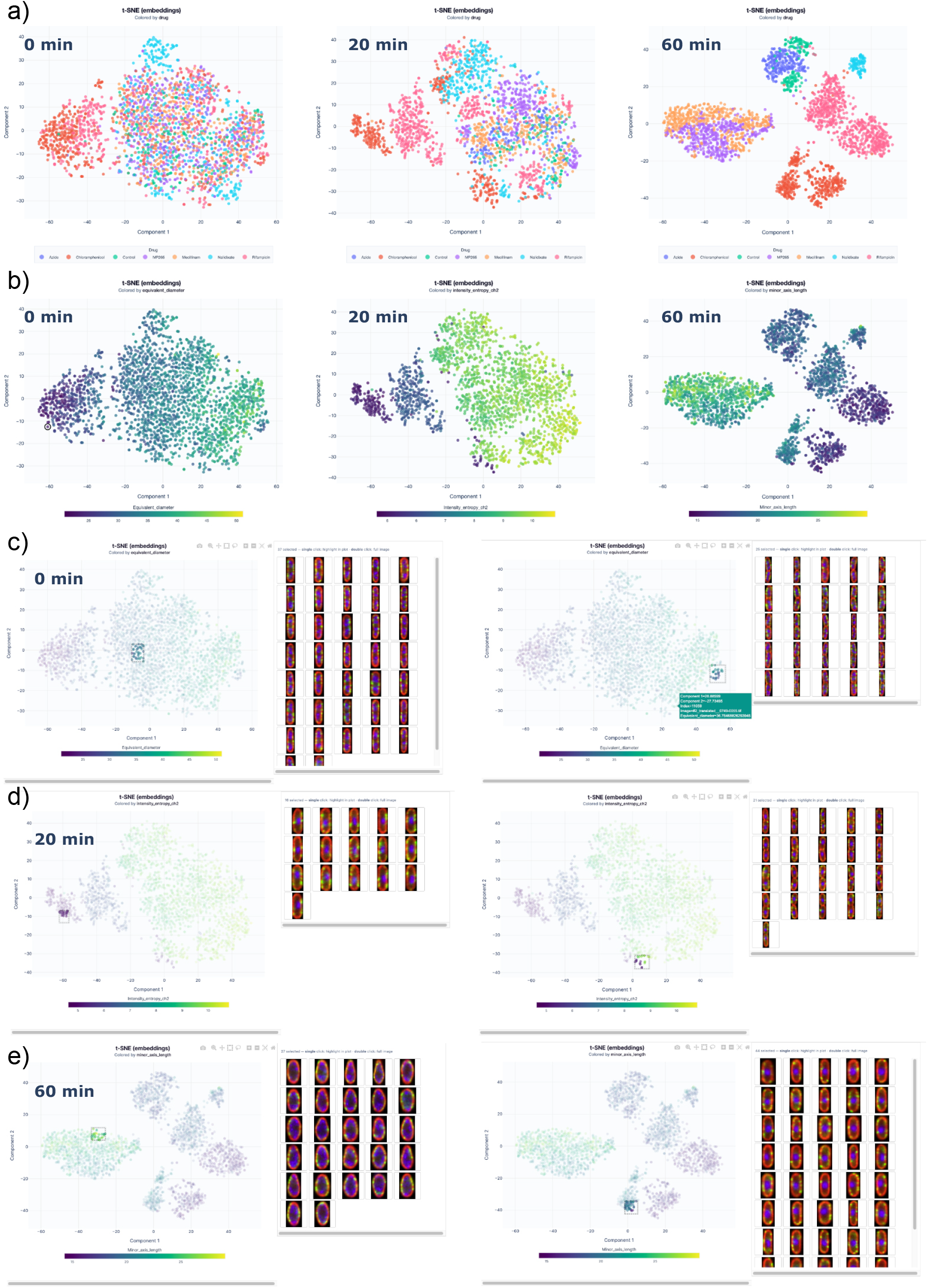
*E. coli* embeddings over time. t-SNE projections of DINOv2 ViT-g/14 embeddings at 0, 20 and 60 min time points, a) coloured by drug or b) by the feature with the highest Pearson correlation with the first t-SNE component (see Fig. 5b) — from left to right, equivalent diameter, DAPI channel entropy and minor-axis length. c-e) Screenshots of PhenoMe’s interactive selection at 0, 20 and 60 min time points.

Finally, we illustrate how PhenoMe can be used to study dynamic biological processes and use the embedding space to represent trajectories with predictive potential. We analysed the HeLa chromatin and microtubule time-lapse dataset from the CellCognition Project (37). DINOv2 embeddings of phase-labelled chromatin crops formed a continuous cellcycle-associated trajectory, with apoptotic observations comparatively separated (**Fig. 5e**), physiologically aligned with apoptosis being itself a cellular process occurring potentially at any stage of the cell cycle. Chromatin area and eccentricity, directly relatable to the nuclear shape, were strongly aligned with this layout. For the full time lapse, we used Cellpose-SAM (38) and TrackMate (39) to obtain single-cell crops and tracks. Clustering produced four groups rather than the six annotated cell-cycle groups represented in the tracked subset; segmentation errors and crops containing signals from touching cells, as well as imbalanced representation of each cell cycle phase, probably contributed to this difference. We used bounding-box crops rather than mask-shaped crops to reduce direct segmentation-shape bias. Projecting individual tracks in PhenoMe exposed progression from interphase through anaphase and back to interphase after division (**Fig. 5f-h)**. Together, these time-resolved analyses illustrate how PhenoMe supports extending the study of static groups to continuous processes by transforming the embedding structure into quantitative phenotypic trajectories. This allows contextualising continuous biological variations and enables individual cells to be followed through successive identified states, as for instance cell-cycle-associated morphological states.

## Discussion

Here, we introduced PhenoMe, a computational framework that combines the representational power of pretrained AI models with the interpretability and analytical interactivity needed to explore biomedical image data and guide sub-sequent analyses. PhenoMe brings learnt representations, quantitative image properties, experimental metadata and image retrieval into a common workflow. By linking AI-derived embeddings to complementary measurements and supporting both interactive and programmatic use, it enables users to identify candidate patterns, inspect the images underlying those patterns, export results for downstream analysis and develop more advanced analytical pipelines. The case studies span distinct biological questions, including cell-type organisation, synaptic structure, compound profiling, image-level quality control, antibiotic response and cell-cycle progression. Together, they demonstrate the scope of representation learning and of the PhenoMe workflow.

The choice of vision backbone can substantially affect the organisation of the resulting representation space. Comparisons using the HPA, Neural Activity States and Synaptic Protein Zoo datasets showed that models with different architectures and training data reorganised the same image collections in different ways. PhenoMe makes such differences accessible for inspection and comparison, which consequently, can support the detection of biological relationships. This was particularly evident in the neural synaptic protein clusters datasets. In the Neural Activity States dataset, STED-FM exposed a treatment-associated organisation that was substantially less apparent in the DINOv2 representation. Likewise, in the Synaptic Protein Zoo it produced a distinct RIM1/2-enriched region and more coherent organisation of the remaining synaptic proteins. By allowing alternative representations to be compared, linked to experimental metadata and inspected through retrieval of the corresponding images, PhenoMe helps identifying a representation in which different conditions or phenotypes could be more readily resolved. These findings illustrate a discovery-enabling role for PhenoMe: the framework can reveal candidate biological relationships and nominate images or groups for targeted quantification and experimental validation. They do not, however, remove the need to validate the selected model and the resulting hypotheses using appropriate experimental controls, quantitative performance metrics and expert assessment.

The joint analysis of learnt and classical features is central to this approach. In BBBC014v1, variation in the FITC signal helped to interpret the embedding gradient associated with TNF-*α* treatment. In the *E. coli* data, minor-axis length linked the organisation of the representation space to antibiotic-associated changes in bacterial width and rounding. In CellCognition, chromatin area and eccentricity varied along the representation-space structure associated with cell-cycle progression. These associations do not establish biological causality. However, when combined with retrieval of the corresponding images, they help users assess whether a pattern is consistent with plausible biological variation or is more likely to reflect a technical artefact, and formulate hypotheses for confirmatory analysis. Likewise, quantities such as local neighbourhood composition, distances between phenotypic groups and trajectories through the representation space can summarise relationships among biological states without requiring every observation to be assigned immediately to a discrete class.

The biological resolution of such analyses depends on the available inputs. Whole images can be embedded without masks or extensive annotations, making broad exploration possible with limited preprocessing. By contrast, objectlevel morphology, single-cell crops and temporal trajectories provide more specific biological context, but require prior segmentation and, for longitudinal analyses, reliable tracking. The *E. coli* and CellCognition case studies illustrate these complementary levels of analysis. Importantly, these two examples also show that AI-derived embedding spaces can represent continuous biological change, rather than only separate static phenotypes. In the *E. coli* data, the temporal ordering of images across antibiotic conditions, together with the associated increase in cell width and rounding, describes a population-level phenotypic response progression. In CellCognition, where single-cell and temporal information are available, successive observations can instead be followed through the embedding space to describe cell-cycle progression and transitions between morphological states. Embedding-space analyses can therefore provide quantitative coordinates in which the direction, rate, continuity or divergence of a biological response may be examined.

This representation of dynamic processes creates opportunities for predictive analysis as described previously (19). For example, the position and recent trajectory of a cell in the embedding space, potentially combined with morphology and experimental metadata, could be used as input to downstream models that predict a subsequent response, state transition or cell fate. PhenoMe enables such candidate predictive features and trajectories to be identified and exported for model development.

As new general-purpose and microscopy-specific models emerge, PhenoMe’s common model interface will allow them to be incorporated and compared within the same analytical workflow. PhenoMe contributes to the broader shift towards analysing microscopy data in learnt representation spaces, whose interpretation depends on connecting abstract image features to visual evidence, experimental context and measurable biological properties. Our results also highlight opportunities to develop methods for comparing datasets, integrating multimodal information and characterising cellular behaviour through the structure and dynamics of embedding spaces. By enabling quantitative inspection, visual validation, export and downstream analysis of learnt representations, PhenoMe provides foundational infrastructure for integrating advances in AI and computer vision into microscopydriven biological discovery.

## Data availability

All datasets used in this study are publicly available. The *E. coli* antibiotic perturbation dataset is from Spahn et al. (36). The CellCognition cell division dataset is from Held et al. (37). The BBBC014v1 (Cytoplasm–Nucleus Translocation) and BBBC021v1 (Compound Profiling) benchmark datasets are from the Broad Bioimage Benchmark Collection (27, 32) and are available at https://bbbc.broadinstitute.org (BBBC014v1, BBBC021v1). A ready-to-use version of the Human Protein Atlas immunofluorescence images as provided in (40) was used. All the information on how to obtain the original datasets as well as the complete metadata is available at the official website of the Human Protein Atlas resource (28) (https://www.proteinatlas.org/humanproteome/subcellular). The Neural Activity States and Synaptic Protein Zoo STED images are from Wiesner et al. (30, 31) and were analysed as released with STED-FM (24). The Synaptic Protein Zoo dataset is also available publicly on Zenodo (https://zenodo.org/records/15480470). Both datasets can be downloaded and preprocessed via the provided notebook in PhenoMe’s repository. Further dataset and preprocessing details are given in the Methods. For reproducibility, ready-to-use checkpoints and processed datasets are available at https://zenodo.org/records/21809769.

## Code availability

PhenoMe is publicly available under an open-source license at https://github.com/PhenoMe-Ecosystem/PhenoMe, and the accompanying napari plugin is available at https://github.com/PhenoMe-Ecosystem/PhenoMe-napari-GUI. The package is also installable from PyPI as phenome. Documentation, tutorials, and example workflows are available in the repository. Scripts and documentation to reproduce the experiments and regenerate figures are provided alongside the tutorial notebooks.

## ACKNOWLEDGEMENTS

We thank Anthony Bilodeau, Frédéric Beaupré, and Gabriel Leclerc for insightful preliminary discussions on the integration of STED-FM into PhenoMe. We also thank the maintainers of public datasets that make these methods reproducible and possible.

This work was partially supported by grants PID2024-157485NB-I00, RED2024-153844-T and AIA2025-164165-C42 (to I.A-C.) funded by MICIU/AEI/10.13039 /501100011033 and by FEDER, UE. Publication funded by grants to support the activities of research groups of the Basque university system 2026-2029 (IT2001-26) (to I.A-C.). E.G.M. acknowledges funding by Fundação para a Ciência e a Tecnologia, Portugal (2023.09182.CEECIND/CP2854/CT0004) and the support of NOVA IMSB through the EU Horizon Europe 101060346. Funded by the European Union. However, the views and opinions expressed are those of the authors only and do not necessarily reflect those of the European Union. Neither the European Union nor the granting authority can be held responsible for them. The AI for Health Platform PREDIS is supported by funding from the Ministère de l’Économie, de l’Innovation et de l’Énergie (MEIE) and the Fonds de recherche du Québec – Santé (FRQS). This project was supported by an IVADO Neuro-AI Research Cluster Catalyst Grant (awarded to F.L.C.), funded through the Canada First Research Excellence Fund (CFREF)(to F.L.C.). G.J. acknowledges funding from the Research Council of Finland (338537, 371287, and 374180 to G.J.), the Sigrid Juselius Foundation (to G.J.), the Cancer Society of Finland (Syöpäjärjestöt; to G.J.), and the Solutions for Health strategic funding for Åbo Akademi University (to G.J.). The research was also supported by the InFLAMES Flagships Program of the Research Council of Finland (decision numbers: 337530, 337531, and 357910). G.J. is supported by the Finnish Cancer Institute (K. Albin Johansson Professorship). I.H.C. is funded by the Finnish Doctoral Program Network in Artificial Intelligence, AI-DOC (decision number VN/3137/2024-OKM-6).

## Supplementary material

### A. Pipeline Architecture

1. PhenoMe implements a modular, dual-stream analysis architecture (Figure 1). Raw images traverse two separate pathways: a deep learning stream that extracts high-dimensional embeddings and a classical property stream that calculates biologically interpretable features. The framework supports the flexible selection of vision backbones, handles inference via a common interface, and independently composes downstream analysis modules for property extraction, batch correction, distance computation, and multivariate analysis. To facilitate large-scale analysis and reproducibility, intermediate data and configurations are lazily processed and stored in an efficient, portable HDF5 format (41). This single-file storage architecture simplifies the sharing of entire analysis results and embeddings among researchers, ensuring consistent exploration across different platforms.

Comprehensive documentation, including API references and interactive tutorials, is provided to ensure that all analyses presented in this study are fully reproducible by the community.

### B. Datasets and Preprocessing

We evaluated PhenoMe on seven diverse microscopy datasets:

- ***E. coli* antibiotic screen**: High-resolution confocal dataset of *E. coli* K-12 treated with antibiotics over time (36). Using published regions of interest (ROIs), we generated bounding boxes to crop every single individual bacterium and produced segmentation masks for classical feature extraction. Multichannel single-cell crops (15,792 across 7 treatments and 7 time points) were rotation-aligned to minimize orientation confounds.
- **CellCognition cell division dataset**: HeLa “Kyoto” cells expressing H2B-mCherry (chromatin) (37). To extract singlecell crops from raw videos, we used *cellpose-sam* (38, 42) for instance segmentation and *TrackMate* (39) for temporal tracking as an external preprocessing step. Segmentation masks were obtained using *cellpose-sam* or, when available, masks provided by the original dataset authors. Bounding boxes were generated to extract cells in each frame, with metadata saved to facilitate subsequent analysis in PhenoMe. Segmented sub-crops (811 cells) represent a ground-truth trajectory of mitotic progression across six cell-cycle classes.
- **BBBC014v1 Cytoplasm–Nucleus Translocation**: Dual-channel (DAPI, FITC) images of MCF7 and A549 cells subjected to a 12-point TNF*α* concentration gradient (32). Analysis included 96 full-field images (48 per cell line).
- **BBBC021v1 Compound Profiling**: MCF7 cells exposed to 113 small molecules (32). We analyzed a subset comprising experiments from the first three weeks, filtered for treatments expressing a detectable phenotypic difference (1080 images); Taxol was excluded from this subset due to its high predominance in the early experimental weeks. Ground-truth mechanism of action (MoA) annotations were obtained from Ljosa et al. (27).
- **Human Protein Atlas (HPA) immunofluorescence**: Subcellular immunofluorescence microscopy mapped to annotated localization patterns (28). The full dataset (113,567 images) was embedded with Cell-DINO (1), DINOv2 ViT-S/14 (21), and ConvNeXt Large (29).
- **Neural Activity States (NAS)**: STED images of PSD-95 protein clusters under various experimental conditions (0Mg2+/Gly/Bic, 48h TTX, Glu/Gly) (24, 30), cropped to 224×244 pixels tiles.
- **Synaptic Protein Zoo**: STED images of synaptic protein clusters, annotated for segmentation via a citizen science project on the Zooniverse platform (https://www.zooniverse.org/projects/reber199/synaptic-protein-zoo). Crops of individual clusters were extracted from the segmentation masks.

Standard preprocessing included intensity normalisation and resizing to vision backbone dimensions. All classical property analyses utilized PhenoMe’s `complete` property preset.

### C. Deep Embeddings

PhenoMe is a model-agnostic framework, compatible with any vision architecture that generates a single embedding vector from an input image. While various architectures and pooling strategies can be utilized, for the Vision Transformers (ViT) (43) used in most of our experiments (specifically DINOv2 (21)), we use the LayerNorm-normalized class (cls) token output from the final transformer layer as the global descriptor. For convolutional networks such as ConvNeXt (29), the global descriptor is obtained from the final classification layer.

To handle multichannel fluorescence images, we employed the **Split Mode** strategy for all relevant analyses, independently extracting embeddings from each channel and concatenating them to form the final embedding vector, preserving channel-specific information. Alternatively, PhenoMe supports a **Combined Mode** that keeps the original image and passes all of its channels as a single input before obtaining the embeddings. When a model requires RGB images, a parameter converts the input to three channels in either mode. In Split Mode each channel is repeated. In Combined Mode the conversion depends on the channel count: one channel is repeated, two channels are zero-padded, and channels beyond the third are dropped. In practice, Split Mode is generally recommended when independent channel variation (e.g., independent stainings) is biologically meaningful, whereas Combined Mode offers greater computational efficiency and can be beneficial when analysing spatial cooccurrence or colocalization or when using a specialised model.

### D. Classical Image Properties

Parallel to embedding extraction, the classical property stream computes biologically inter-pretable metrics using the `complete` preset, generating a comprehensive taxonomy of features per channel:

- **Shape descriptors:** Requires a segmentation mask. Extracted via `regionprops` (44), including area, perimeter, major and minor axis lengths, eccentricity, solidity, orientation, equivalent diameter, extent, and Euler number. Derived metrics include circularity.
- **Whole-image intensity:** Computes mean, standard deviation, minimum, and maximum pixel intensity across the entire unmasked image.
- **Masked intensity:** Constrains intensity statistics (mean, std, min, max) to the foreground segmented region.
- **Concentric ring profiles:** Using the segmentation, the object is radially divided into N concentric rings of equal radial width using Euclidean distance transforms from the boundary, where ring 1 represents the surface of the object and ring N represents the core. The mean and standard deviation of intensity are computed per ring to capture spatial gradients.
- **Texture (GLCM):** Gray-Level Co-occurrence Matrix features (contrast, dissimilarity, homogeneity, energy, correlation) computed at a pixel distance offset of *d* = 1 and 4 angles, then averaged, measuring local spatial intensity patterns.
- **Image quality metrics:** Focus quality is estimated via a multi-scale blur effect metric (`sharpness_metric`), along-side `intensity_entropy` (Shannon entropy) to quantify overall structural complexity.

For multichannel images, intensity, texture, and quality properties are computed and suffixed independently for each channel.

### E. Batch Correction

In this manuscript, we analysed the raw embeddings to examine the organisation produced by off-the-shelf models. However, uncorrected representations may be inconvenient in some cases: variation between plates, acquisition days, instruments or sample-preparation protocols may obscure the signal of interest. To mitigate such technical variation across experimental batches, PhenoMe integrates two common correction strategies applicable to either the embedding or classical property streams: **Standardization** (Z-scoring) and **Sphering** (ZCA Whitening) (45). Both methods adjust sample features relative to reference control statistics from the same batch.

### F. Dimensionality Reduction and Clustering

High-dimensional data can be visualized and analyzed using standard dimensionality reduction techniques: Principal Component Analysis (PCA) (34), t-SNE (35) and UMAP (46). Unsupervised grouping can be performed using *k*-means, DBSCAN (47), and Gaussian Mixture Models (GMM) (45). To ensure exact reproducibility across runs, all stochastic algorithms were initialized with a fixed global random seed.

### G. Phenotypic Distance Quantification

Phenotypic magnitude relative to a reference state (e.g., negative control, time zero, or any chosen target class) is quantified using mathematical distance metrics in the embedding or property space. PhenoMe computes distances in two modes:

- **Centroid distance:** Distance from a query point to the mean vector of the reference class.
- **Nearest-neighbor distance:** The minimum distance from a query point to any other sample in the reference class. The distance metric employed is either Euclidean distance or Cosine distance (1 *−* cosine similarity).

### H. Interpretability via Feature Correlation

PhenoMe bridges the gap between deep representation and physical space by statistically linking the two streams. It operates in two explanatory modes:

- **Component Correlation:** Associates derived embedding coordinate axes (e.g., PCA or t-SNE dimensions) with classical property vectors.
- **Embedding–Property Correlation:** Maps raw, individual embedding dimensions directly to physical traits.

Relationships are evaluated pairwise using Pearson correlation (34), Spearman rank correlation (48), distance correlation (49), or mutual information (50). For multivariate interpretation of a single embedding axis, PhenoMe applies **LASSO** (51) or **Random Forest** regression (52). This isolates a sparse, compact subset of classical features that linearly or non-linearly predicts variation along the embedding axis, offering an interpretable description of high-dimensional structure.

### I. Statistical Analysis

Statistical associations (Pearson *r*, Spearman *ρ*) and multivariate scores were computed using scikit-learn (53). To flag extreme phenotypes and identify technical acquisition anomalies (e.g., staining artifacts), outliers were isolated from the distance to the group centroid in embedding space, using a z-score or interquartile-range threshold. Property enrichment analysis uses leave-one-group-out z-scores to highlight defining characteristics of clusters. Fixed random seeds were used to ensure reproducible sampling and statistical evaluation across all experiments.

#### I.1 Environment and Reproducibility

PhenoMe can be installed as a standard Python package via pip or using provided conda environments for both CPU and GPU setups. The analyses presented in this study were performed using Python 3.12 and PyTorch 2.12 (54). As an open-source tool, PhenoMe is actively maintained; for the most up-to-date system requirements, installation instructions, and framework compatibility, users should refer to the official documentation and repository. Reproducibility is enforced via `PhenoMe(seed=42)`, which fixes random seeds for Python, NumPy, PyTorch, CUDA, clustering, and dimensionality reduction. Scripts and instructions to reproduce the experiments are available in the repository, alongside tutorial notebooks for regenerating the main analyses.

#### I.2 Interactive Tutorials and Documentation

To lower the barrier to entry and facilitate rapid adoption, PhenoMe is accompanied by a comprehensive suite of interactive tutorials. These tutorials are provided as Jupyter notebooks and cover the entire workflow, from data ingestion and model wrapping to batch correction and interpretability analysis. Furthermore, the official documentation provides detailed API references, implementation guides, and best practices for extending the toolkit with custom vision backbones or analysis modules.

### J. Visual outputs

## Footnotes

1 https://www.zooniverse.org/projects/reber199/synaptic-protein-zoo

